# Phasic modulation of visual representations during sustained attention

**DOI:** 10.1101/2020.09.04.282715

**Authors:** Mats W.J. van Es, Tom R. Marshall, Eelke Spaak, Ole Jensen, Jan-Mathijs Schoffelen

## Abstract

Sustained attention has long been thought to benefit perception in a continuous fashion, but recent evidence suggests that it affects perception in a discrete, rhythmic way. Periodic fluctuations in behavioral performance over time, and modulations of behavioral performance by the phase of spontaneous oscillatory brain activity point to an attentional sampling rate in the theta or alpha frequency range. We investigated whether such discrete sampling by attention is reflected in periodic fluctuations in the decodability of visual stimulus orientation from magnetoencephalographic (MEG) brain signals.

In this exploratory study, human subjects attended one of two grating stimuli while MEG was being recorded. We assessed the strength of the visual representation of the attended stimulus using a support vector machine (SVM) to decode the orientation of the grating (clockwise vs. counterclockwise) from the MEG signal. We tested whether decoder performance depended on the theta/alpha phase of local brain activity. While the phase of ongoing activity in visual cortex did not modulate decoding performance, theta/alpha phase of activity in the FEF and parietal cortex, contralateral to the attended stimulus did modulate decoding performance. These findings suggest that phasic modulations of visual stimulus representations in the brain are caused by frequency-specific top-down activity in the fronto-parietal attention network.

## 1. Introduction

The ability to efficiently respond to the environment according to current behavioral goals relies on the process of attention to prioritize information. The brain has a limited processing capacity and therefore, external stimuli and internal mental states compete for cognitive resources, for example by focusing on a specific location in space. According to our subjective experiences, this struggle happens continuously: you can be so focused on a traffic light in order to hit the gas the moment it turns green, that you are totally oblivious to an acquaintance who waves at you from the sidewalk. We seem to attend to stimuli in a continuous fashion, but recent evidence suggests that, instead, our attention waxes and wanes in discrete, periodic cycles.

Early indications of discrete sampling come from psychophysical experiments, showing that there are limitations on the number of temporal events one can perceive per unit of time. It turns out that when visual flashes are presented to a subject at a constant rate, the subject can only account for a maximum of 10-12 items/second, even when the presentation rate is higher (White and Harter, 1969). In another experiment, reaction time distributions were found to be multi-modal, with ∼100 ms in between the peaks, indicating a periodicity of about 10 Hz (Venables, 1960). Given that attention has a facilitating effect on perception (Hawkins et al., 1990), the rhythmicity in reaction times suggests that attention might be discrete and rhythmic.

More recently, neuroscientific experiments have linked the rhythmic modulation of behavior to brain activity, in particular to rhythmic synchronization of cortical excitability in relation to visual attention. For example, Busch et al. (2009) found that the detection of a near-threshold stimulus was related to the phase of spontaneous rhythmic activity in the theta/alpha band (7-10 Hz) just before stimulus onset. In a similar experiment, Busch and VanRullen (2010) later found that this effect was only present when the subject actively attended the near-threshold stimulus. Thus, according to these studies the phase of the theta/alpha rhythm is inherently related to fluctuations in visual attention and, with that, behavior. Helfrich et al. (2018) pinpointed these top-down modulatory effects to the fronto-parietal attention network, specifically the frontal eye fields (FEF), the intraparietal sulcus (IPS), and inferior and superior parietal regions.

The behavioral evidence that suggests that visual perception might inherently be discrete is thus supported by neural evidence that ascribes these effects to the phase of spontaneous activity. However, it remains unclear why visual attention is so strongly related to theta/alpha activity. According to theoretical accounts of visual processing and attention (Jensen et al., 2014; Klimesch, 2012), these rhythms prioritize input by inhibiting neural processing. In short, neuronal firing is inhibited at oscillatory peaks, but once inhibition ramps down (i.e. in the downgoing flank and trough of the oscillation), visual representations will activate according to their excitability. We investigated whether a discrete sampling of attention affects the input gain of incoming stimuli, i.e. whether the strength of a stimulus’ representation in brain activity shows periodic fluctuations.

Developments in the analysis of neuroimaging data have resulted in the ability to decode mental states from non-invasive measurements (Haynes and Rees, 2006). This has been particularly successful in the visual domain in both fMRI and M/EEG (van de Nieuwenhuijzen et al., 2013; Zafar et al., 2015). The high temporal resolution of M/EEG enables us to ask questions about the temporal evolution of visual presentations using multivariate pattern analysis (MVPA; Cichy et al., 2015; Pantazis et al., 2018; Ramkumar et al., 2013). In order to investigate whether the strength of visual representations in brain activity shows periodic fluctuations, in this exploratory study we decoded visual stimulus information from MEG activity as a function of instantaneous phase of the ongoing activity. Human subjects participated in a spatial attention task while MEG was being recorded. The visual stimuli consisted of oriented gratings, with two possible orientations. If the representations of these stimuli are activated only in distinct time windows as a result of discrete attentional sampling, we expect the decoding performance to fluctuate accordingly, with a dependency on the theta/alpha phase. Because early visual cortex is most sensitive to oriented gratings (Hubel and Wiesel, 1962), this modulating signal was expected to be present there. Alternatively, the modulatory signal could come from the fronto-parietal attention network, given its putative role in attention (Buschman and Kastner, 2015) and evidence that points to this network in the phasic modulation of behavior (Busch et al., 2009; Helfrich et al., 2018).

## 2. Methods

### 2.1 Subjects

10 healthy volunteers participated in this study, of which 3 male and 7 female. Their age range was 19-27 (mean ± SD: 24 ± 2.3). All subjects had normal or corrected-to-normal vision, and all gave written informed consent according to the declaration of Helsinki. This study was approved by the local ethics committee (CMO region Arnhem/Nijmegen) and conformed to the Declaration of Helsinki.

### 2.2 Experimental design

#### 2.2.1 Stimuli

The experimental task was programmed in MATLAB (R2013a, Mathworks, RRID: SCR_001622) using Psychophysics Toolbox (Brainard and Vision, 1997), RRID: SCR_002881). All stimuli were presented against a black background (figure 1). A fixation cross (1.8 visual degrees (°)) was presented for a baseline period (800 ms), after which one of the horizontal arms widened to act as spatial cue (800 ms). Then two sinusoidal gratings (3.2°) appeared on the lower part of the screen, one in either hemifield. The gratings were drifting upwards (1.33 cycle per degree, drift rate 1 Hz) and could be oriented clockwise (CW, 45°) or counter-clockwise (CCW, 135°). After a variable stimulus duration (0.5-1.5 s) one of the gratings rotated clockwise or counterclockwise with a variable rotation angle. The gratings then remained drifting for 100 ms, after which a question mark was presented to indicate the response window (max. 1200 ms).

**Figure 1.**
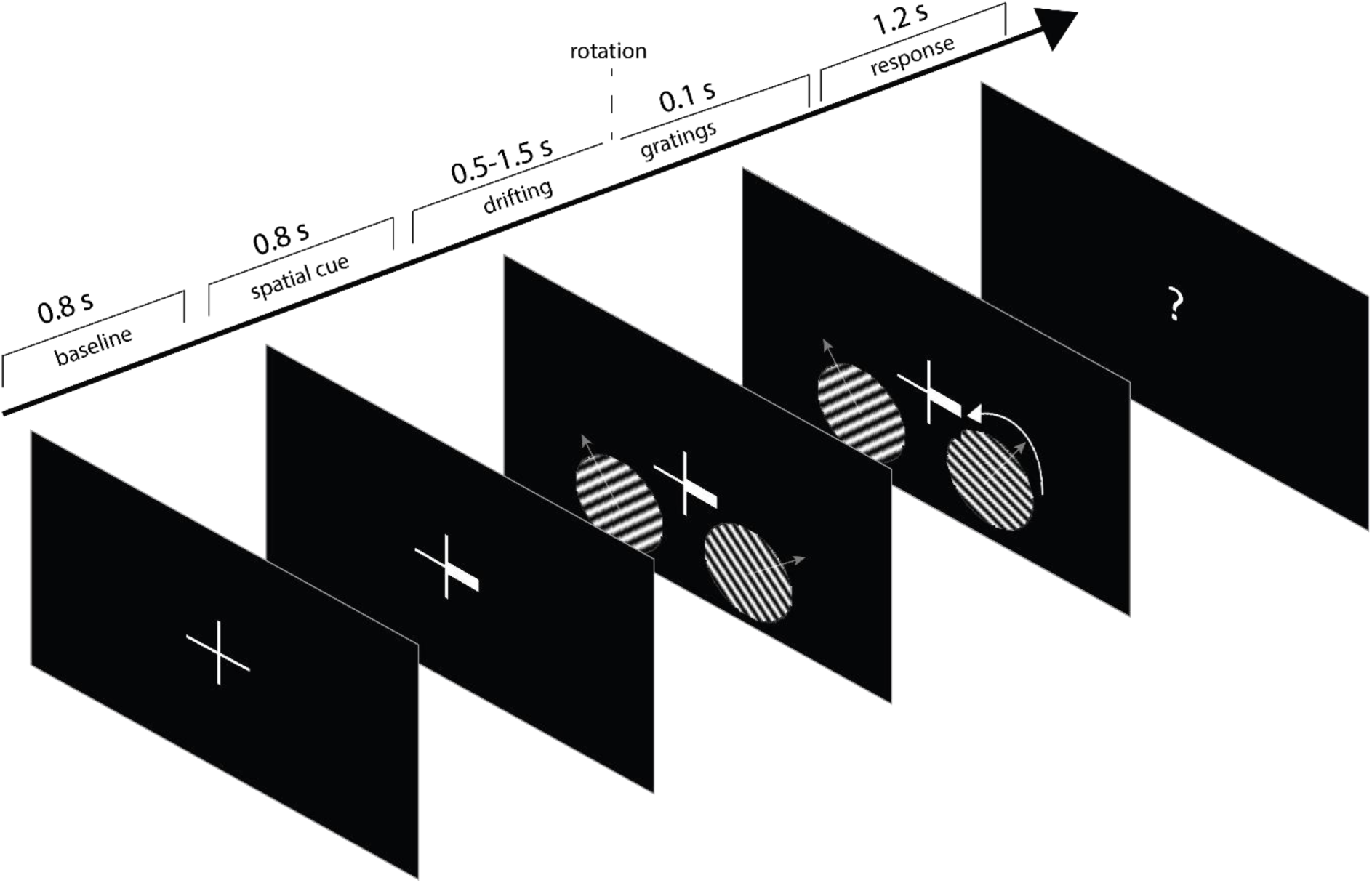
Trial timeline. Subjects had to look at the center of a fixation cross throughout the trial. After a baseline period, a spatial cue appeared, indicating the target hemifield for covert visuospatial attention. Then two oriented, drifting gratings were presented, after which the rotation direction of one of them had to be indicated with a button press. Semi-transparent arrows in the gratings indicate direction of movement of the drifting grating.

#### 2.2.2 Experimental equipment

Stimuli were presented by back-projection onto a semi translucent screen (width 48 cm) by an Eiki LC-XL100L projector with a refresh rate of 60 Hz and a resolution of 1024 ⨯768 pixels. Subjects were seated at a distance of ∼76 cm from the projection screen in a magnetically shielded room. MEG was recorded throughout the experiment with a 275-channel axial gradiometer CTF MEG system at a sampling rate of 1200 Hz. In addition, subject’s gaze direction and pupil size were continuously recorded using an SR Research Eyelink 1000 eye-tracking device (RRID: SCR_009602) at 1000 Hz and resampled to 1200 Hz. This resulted in three extra channels that were saved together with the MEG data: the horizontal and vertical gaze position and pupil diameter.

Head position was monitored in real-time during the experiment by using head-positioning coils at the nasion and left and right ear canals of the subject (Stolk et al., 2013). Subjects were adjusted to the starting position of the first session at the start of a next session. Additionally, subjects readjusted to the original position when the head position deviated more than 5 mm from it. Behavioral responses during the MEG session were recorded using a fiber optic response pad (FORP). In addition to the MEG recording, anatomical T1 scans of the brain were acquired with one of the 1.5/3 T Siemens MRI systems present at the facility (Siemens, Erlangen, Germany). In order for co-registration of the MEG and MRI datasets, the scalp surface was mapped using a Polhemus 3D electromagnetic tracking device (Polhemus, Colchester, Vermont, USA).

#### 2.2.3 Procedure

The experiment contained three 1-hour MEG sessions per subject. Each session consisted of a maximum of 10 blocks of 80 trials or until the end of the session, with a self-determined break in between the blocks. Subjects were instructed to keep their head as still as possible and, preferably, to blink at the end of every trial (see figure 1 for trial timeline). Subjects had to fixate at the fixation cross at all times and pay covert attention to the cued grating, instructed by the fat arm of the fixation cue. Both gratings could either have a clockwise or counterclockwise orientation and were presented for either 0.5 (10%), 1.0 (80%), or 1.5 (10%) seconds. Only the 1s trials were of interest, the other timings functioned to exclude the possibility that the subject would adapt their attention only after a fixed period following the stimulus onset. Then one of the gratings (cued grating in 80% of 1-second trials) rotated either clockwise or counterclockwise. The amount of rotation was adjusted online throughout the experiment such that the performance level would be at 80% for all validly cued, 1-second trials, and was initiated at 1.5°. Subjects had to indicate the rotation direction with a button press of the index finger (CW: right index finger; CCW: left index finger). If the subject was too slow (i.e. slower than 1200 ms) the question mark in the response window turned red for 100 ms, and a new trial was initiated. All trials were counterbalanced regarding grating orientation, cue validity, and stimulus presentation time, and trial order was randomly permuted per session.

### 2.3 Data analysis

All analyses of MEG data were executed in MATLAB (2018b, Mathworks, RRID: SCR_001622) using the FieldTrip toolbox (Oostenveld et al., 2011) and custom-written code. Anatomical T1-scans were processed with FreeSurfer (RRID: SCR_001847) and HCP’s Connectome Workbench (RRID: SCR_008750), and MATLAB.

#### 2.3.1 MEG preprocessing

MEG data were preprocessed for each session separately. First, excessively noisy channels and trials were removed from the data by visual inspection, including trials containing SQUID jumps and muscle artifacts. Similarly, eye-tracking data were inspected to remove trials containing eye blinks or saccades in the period [-1, 1] seconds relative to stimulus onset. The data were then demeaned and high pass filtered at 0.1 Hz using a finite impulse response windowed sinc (FIRWS; Widmann, 2006) filter. Additionally, line interference at 50 Hz was removed using a discrete Fourier transform (DFT) filter, as well as its harmonics at 100 and 150 Hz. Signals related to cardiac activity or eye movements were manually identified and removed using independent component analysis (ICA) and visual inspection. This was done prior to the removal of trials containing residual eye movements and blinks. Finally, the data were down sampled to 200 Hz. For all further analyses only trials were used with a valid cue, a correct response, and of a duration of 1 second (stimulus onset to stimulus change).

#### 2.3.2 MRI preprocessing

MRI data were co-registered to the CTF coordinate system using the head-positioning coils and the digitized scalp surface. Volume conduction models of the head were created using a segmentation of the anatomical image, with SPM8 (Penny et al., 2011). Freesurfer and Connectome Workbench were used to construct source models, with dipole positions positioned on a cortically constrained surface, containing 15,684 dipole locations. Subsequently, forward models were computed from individuals’ structural MR images using a single-shell volume conduction model.

#### 2.3.3 Time-frequency analysis

Time-frequency resolved power was estimated for initial exploration of the data. This was done separately for low (2-30 Hz) and high (30-80 Hz) frequencies. First, the data were transformed to synthetic planar gradients (Bastiaansen and Knösche, 2000) and padded to 4 s. For low frequencies, a Hanning tapered sliding time window of 500 ms was slid over these data in steps of 100 ms, from -1.0 to 1.0 s relative to stimulus onset, resulting in a frequency resolution of 2 Hz. For high frequencies, DPSS multi-tapers were used with a sliding time window equal to five cycles for each frequency and 50 ms steps, 4 Hz frequency resolution and frequency specific smoothing (20 % of the frequency value). After spectral decomposition synthetic planar gradients were averaged to form a single spectrum per sensor.

High frequency time-frequency spectra are plotted relative to the average power in the baseline. Low frequency time-frequency spectra are plotted as the Attentional Modulation Index (AMI), the relative increase in attend left versus attend right trials:

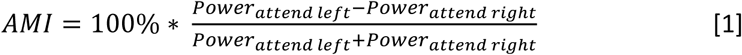

#### 2.3.4 Virtual channel of induced gamma band activity

In order to bin data according to phase, a phase-providing signal is required. Since gamma power is closely related to active stimulus processing, the location in the brain with the maximum gamma power increase can be used as a central location for stimulus processing. In order to estimate this location for each subject individually, first the peak gamma frequency was estimated.

A 600 ms window, padded to 1 s, was used to estimate the peak frequency; right before stimulus onset for the baseline estimate, and from 400-1000 ms after stimulus onset for the stimulus-induced estimate (excluding the ERF in the first 400 ms). Power was estimated using a fast Fourier transform (FFT) and a Hanning taper between 30 and 100 Hz, with a frequency resolution of 1 Hz. The relative power increase from baseline was manually inspected in order to select the peak frequency and bandwidth, based on the (posterior) channels with the largest increase.

Another FFT (on the 1 s data) on the subject’s individual gamma peak frequency was used to inform a Dynamic Imaging of Coherent Sources (DICS) beamformer (Gross et al., 2001). The amount of smoothing was tailored to the individual’s bandwidth using DPSS tapers. Spatial filters were created for each of the dipole locations in a 3-dimensional grid, using the cross-spectral density (CSD) estimated from the baseline and stimulus data concatenated, and a regularization of 100% of the mean sensor level power. This was done separately for each channel, after which single trial power in source space was concatenated over sessions. Next, the two dipole locations were selected, one for each hemisphere, that showed the largest increase in gamma power relative to baseline (i.e. the dipole location with the highest T-value resulting from a T-test). Spatial filters for the time domain were estimated with a Linearly Constrained Minimum Variance (LCMV) beamformer for these two locations.

First, the time domain data were baseline corrected based on the 100 ms before stimulus onset. The data from 100 ms before to 1000 ms after stimulus onset were used to estimate the covariance, and spatial filters were created using a regularization of 100 % of sensor-level power.

#### 2.3.5 Brain-wide cortically constrained anatomical parcels

Source modelling can be used to increase the signal to noise ratio of a particular signal of interest and improve spatial consistency over subjects. Without any dimensionality reduction, this can lead to an inefficiently large search space. Therefore, brain-wide time domain data were first modelled on cortically constrained meshes of dipole positions using an LCMV beamformer, and then grouped in 374 parcels based on an anatomical atlas (Conte 69 atlas, Van Essen et al., 2012), as described in van Es & Schoffelen (2019). This was done separately for each session and concatenated afterwards.

#### 2.3.6 Phase estimation

Phase time courses were estimated from either the virtual channel based on the location of maximum gamma power increase (see 2.3.4) or the anatomically defined cortical parcels. The epoched data were padded to 4 s and for each frequency-of-interest a Hanning tapered sliding time window was slid over the data with steps of 5 ms. The time window was equal to two cycles of the frequency-of-interest. The phase was estimated by taking the angle of the Fourier coefficients.

#### 2.3.7 Decoding of stimulus orientation

All decoding analyses were done separately for the attend-left, and attend-right conditions. The goal of the decoding analysis was to decode the orientation of the attended grating (clockwise or counter clockwise) from the MEG or eye tracker data. First, the data were selected from 400 to 1000 ms after stimulus onset. The first 400 ms after stimulus onset were discarded in order not to be biased by superior decoding during the stimulus onset response. All time points of interest (i.e. belonging to a particular phase bin, see 2.3.8) were selected and concatenated, such that each time point functioned as an observation. In order to increase SNR and assuming that the representation of a particular orientation can be generalized over observations, groups of 5 randomly chosen observations (i.e. belonging to the same phase bin) without replacement were averaged together to create pseudo-trials (Guggenmos et al., 2018). The data were pre-whitened and reduced in dimensionality, by applying singular value decomposition (SVD) to the demeaned data, and multiplying the channel-level data with those vectors that explain at least 99 % of the variance in the data. The trials with different orientations (clockwise/counterclockwise) were then split up and equalized in terms of number of observations. After this preprocessing, a support vector machine (SVM) with a linear kernel and 5-fold cross-validation was used to distinguish between the two stimulus orientations. Note that each observation (i.e. trial-time point) was only used for a single pseudo-trial and in only one fold.

In order to assess whether we could decode stimulus orientation with above chance accuracy, this approach was applied to all data and repeated 100 times, where every repetition different subsets of observations were averaged. For an estimate of the null distribution of accuracies, condition labels were shuffled before applying the SVM classifier.

In case of assessing whether we could decode orientation from the MEG data, all data were assigned to the same (phase) bin. In case of assessing phasic modulations in decoding performance, the approach above was repeated for each of 18 phase bins independently. Within a phase bin, the SVM was repeated 20 times, where every time different subsets of observations were averaged. The overall decoding performance was the average accuracy over folds and trial-averaging repetitions. A null distribution was estimated by circularly shifting the phase time courses of every trial at random from anything between -2π and 2π before phase binning the data. This way, the phase binning of the raw data was randomized without breaking the temporal (autocorrelation) structure within a trial. This was repeated 100 times to create a null distribution.

A similar approach was used in a control analysis that tested whether above chance MEG decoding was still possible when taking differences in gaze into account. The data were preprocessed as described above and the channels from the eye tracker were subsequently used in a 10-fold cross-validation. From the training data in each fold, the one trial in each class that was most informative in training the SVM was removed, after which another decoding took place. These steps were repeated until orientation decoding based on eye tracker data was at chance level (evaluated with a T-test over 10 folds versus scrambled class labels). When performance was at chance level, the same trials (i.e. those that were most discriminative in the eye tracker data) were removed from the MEG data. The MEG data were then used in a subsequent decoding, and the accuracy was tested against the randomized version with a T-test over 10 folds.

#### 2.3.8 Phasic modulation of decoding performance

The dependency of decoding performance on the phase of a particular frequency was taken as a measure for the strength of periodic fluctuations of visual representations. For this, either the virtual channels based on maximum gamma power increase were used to provide the phase time course, or the individual anatomical parcels were used. Based on these phase time courses, each data point was assigned to any of 18 equidistant phase bins. The decoding approach described above was applied independently for each bin, resulting in 18 accuracy scores. To this, a cosine function of one cycle was fitted, with amplitude and phase as the two free parameters, estimating the strength of the phasic dependence and the optimal decoding phase. This procedure was repeated for every frequency of interest, each time using the phase time course of one particular frequency to bin the data.

#### 2.3.9 Phasic modulation of behavioral performance

We assessed whether behavioral performance, as quantified by the reaction time, depended on the phase of a particular frequency at the moment of the stimulus change (to which the subjects had to respond) occurred. For all 18 phase bins, the reaction times of all trials with a phase within a quarter cycle distance to the center phase of that bin were averaged. Finally, a cosine function was fitted to the set of average reaction time as a function of phase bin, of which the amplitude indicated the strength of the phasic modulation. This was done for every frequency of interest, and using every anatomical parcel’s time course as a phase-providing signal. In order to create a null distribution of cosine fit amplitude scores, the same approach was repeated 100 times after shuffling the reaction time over trials.

### 2.4 Statistical analysis

#### 2.4.1 Decoding of stimulus orientation

The average decoding accuracy over 100 permutations for observed data (i.e. in which in every permutation different subsets of trials were averaged to increase SNR) was compared with the average decoding accuracy of 100 permutations including randomization of class labels. These were subjected to a dependent samples T-test at the group level, with an alpha of 0.05, corrected for the number of contrasts (attend left trials, attend right trials, decode attended stimulus, decode unattended stimulus).

#### 2.4.2 Phasic modulation

The amplitude of the cosine fit for observed data and for the 100 first-level random permutations (i.e. in which phase-binning was randomized) were subjected to second-level non-parametric permutation tests with clustering across frequencies at the group level (based on 1000 permutations; Maris and Oostenveld, 2007). In short, a null distribution of amplitudes was created for every frequency by averaging one sample from the first-level random permutation distribution over subjects. From this distribution a threshold is established based on the critical alpha level (0.05). The test statistic (i.e. cosine-fit amplitude) of neighboring frequencies that exceed the threshold are clustered and their statistic is summed. This is done both for the observed data and for all random permutations. A p-value is derived by comparing the largest summed statistic from the observed and random data, and counting the occasions in which the observed test statistic was larger than the random statistic. In case of testing phasic modulation on brain-parcels, statistics were constricted to a total of 36 parcels, spanning the left and right FEF and parietal cortex (figure 2).

**Figure 2.**
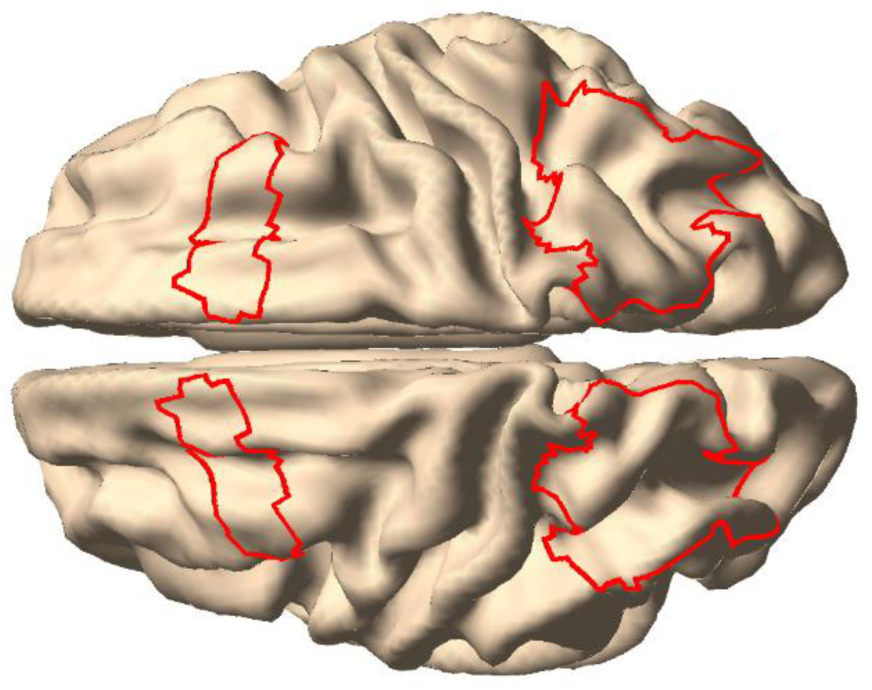
The regions of interest for the phasic modulation of decoding performance and reaction times included parcels in the left and right FEF and parietal cortex.

## 3. Results

10 human participants were cued to covertly attend to one of the two gratings presented in each hemifield, while fixating on a cross in the middle of the projection screen (Figure1). The subjects had to indicate the direction of change of the grating’s orientation with a button press. The number of completed 1-second trials was 1956 on average (SD =18.8). One session from one subject was excluded in its entirety because the behavioral performance was at chance level (i.e. 50 %). Invalidly cued trials and trials containing artifacts were excluded, and in all remaining trials (mean ± SD = 1420 ± 207) the behavioral performance was on average 87% (SD = 3.6 %). The behavioral performance in one session of another subject was inflated (∼95 %) because of an incorrect setting in the experimental procedure. This session is still included and is not believed to have had a profound effect on the results. Excluding incorrect trials, 1235 (SD = 186) trials were considered for further analysis.

### 3.1 Stimulus-induced power changes

In order to assess whether the experimental manipulation led to expected changes in brain activity, we conducted a time-frequency analysis of power in channel space. In the low frequency range (1-30 Hz), we expected differences in posterior alpha power (8-13 Hz) induced by the spatial attention cue. Since alpha power is a proxy for the amount of inhibition in a brain region, it is expected to increase in the hemisphere ipsilateral to the attended hemifield, and to decrease contralateral to the attended hemifield. Figure 3 shows this as the attentional modulation index (AMI, i.e. the relative power difference between attend-left and attend-right trials), with B showing the topography of the alpha band, pre-stimulus onset AMI, and A and C showing the AMI over selected left/right posterior channels on the left and right panels, respectively. Indeed, during presentation of the spatial cue (i.e. before onset of the grating stimulus), there is a relative increase in ipsilateral alpha power with the respect to the cued hemifield and a decrease in contralateral alpha power.

**Figure 3:**
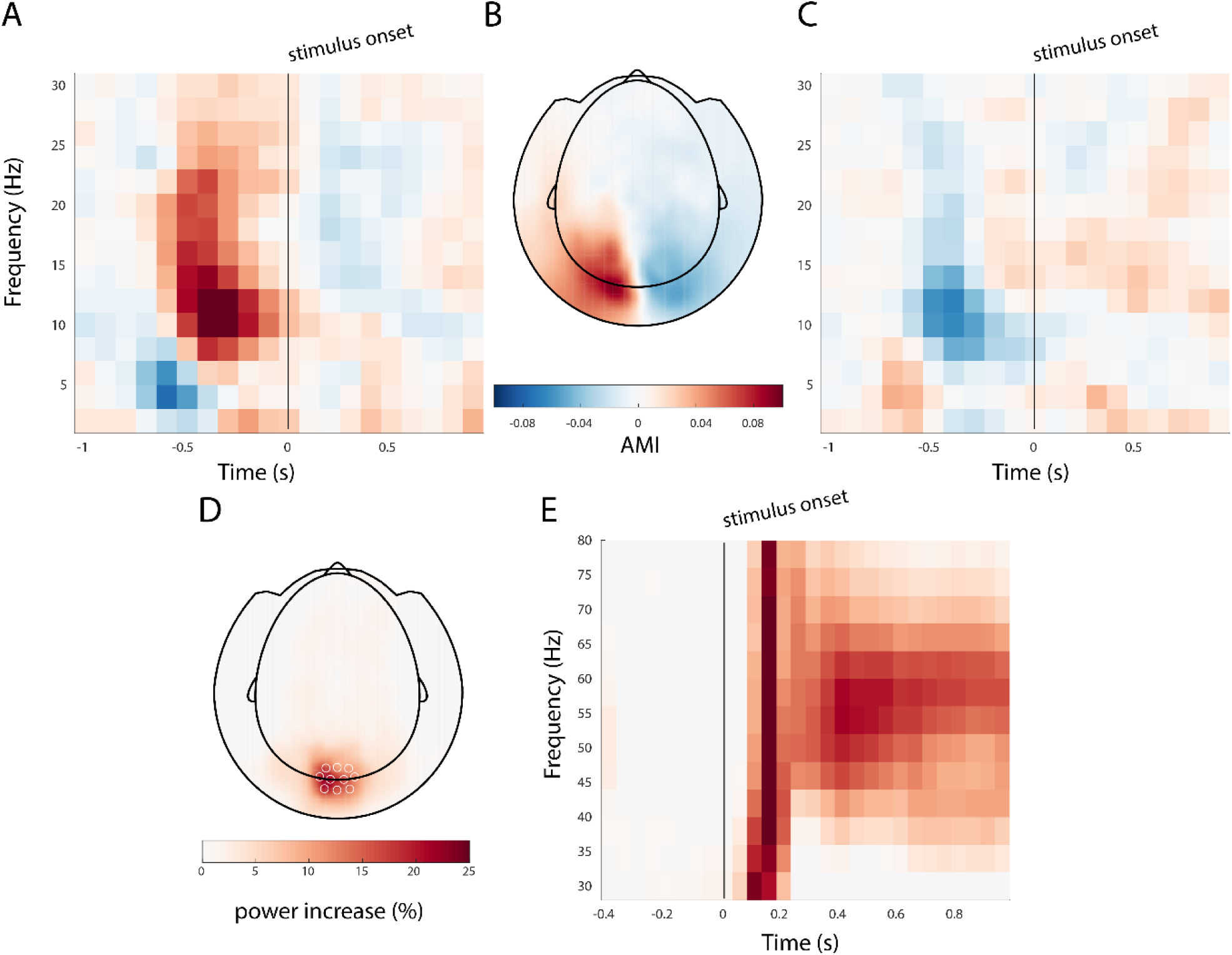
Stimulus induced changes. A, C) Spatial cue (presented at -0.8 ms) induced attentional modulation in posterior alpha power, as indicated by the attentional modulation index (AMI). The AMI over frequency and time was averaged over selected left and right posterior MEG sensors, respectively. B) The topography of the AMI in the alpha band, in the pre-stimulus onset window. D) The topography of the stimulus induced gamma response; E) the time-frequency response, averaged over selected occipital MEG channels.

In the high frequency range (30-90 Hz), we observed an increase in band-limited gamma power, induced by the grating stimuli (Figure 3): a sharp rise in gamma power right after stimulus onset in occipital channels (up to 44 ± 45 %; mean ± SD, figure 3D), with a sustained power increase of 15 ± 14 % (mean ± SD) after 200 ms in the 45-65 Hz band (figure 3E). Source modelling confirmed that the induced gamma power increase originated in posterior brain areas, and the maximum induced gamma power was in occipital cortex in all subjects (figure 4).

**Figure 4:**
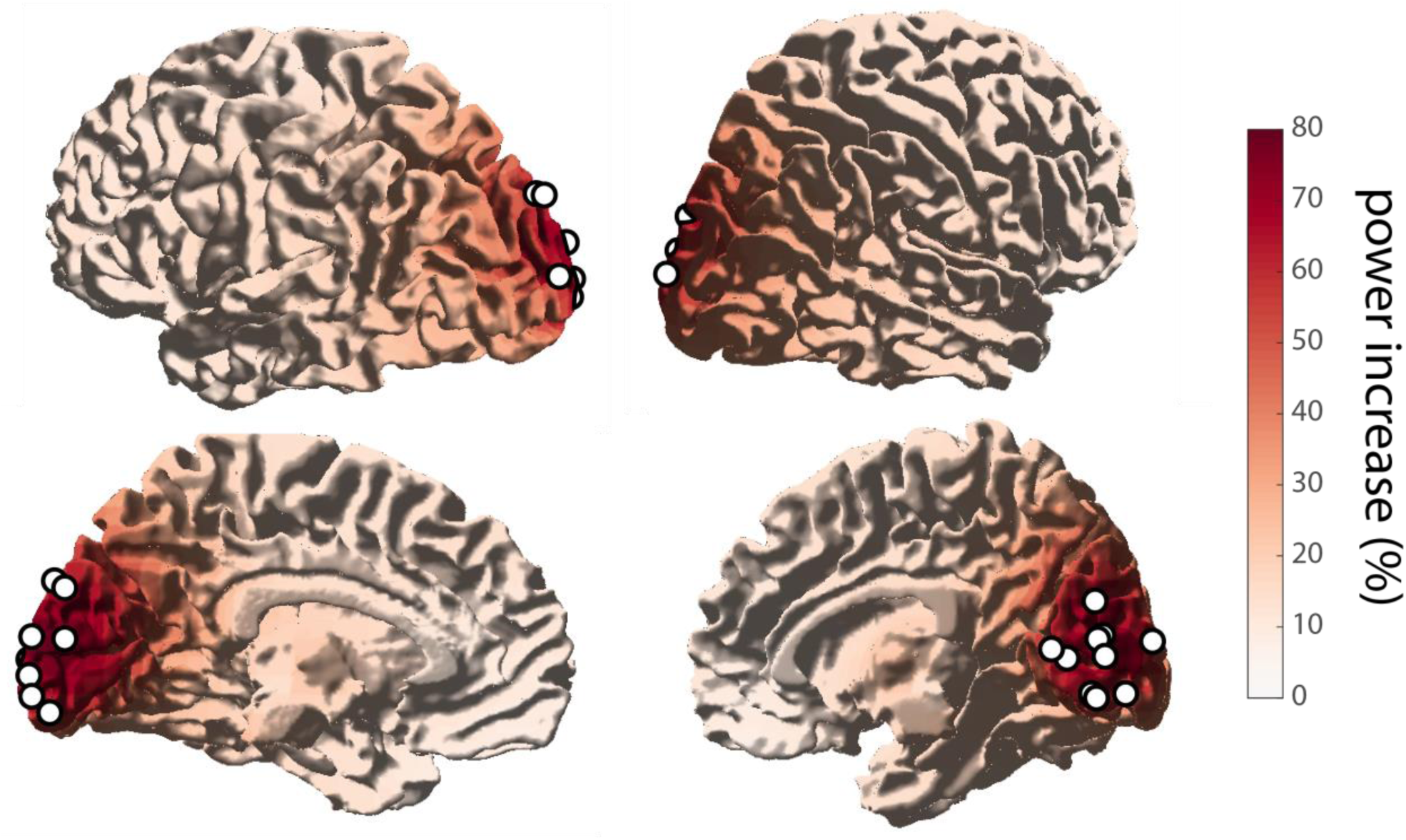
Source of induced gamma activity. Stimulus induced gamma power increase originated from occipital areas, mostly in V1-V2, and close to the midline. Maximum power increase for all subjects are denoted by white circles.

### 3.2 Stimulus orientation can be decoded from the MEG signal

The accuracy of decoding stimulus information from the MEG signal is known to be largest in the 0 to 250-400 ms interval; i.e. when the event-related fields are strongest (ERF; Cichy et al., 2015; Pantazis et al., 2018; Ramkumar et al., 2013). This high decodability is most likely due to the stimulus onset locked transients, and therefore could confound our analysis of a possible phase modulation of decoding accuracy. For this reason, all subsequent decoding analyses excluded data in the time window of the main ERF components, and we only used data from interval 400 - 1000 ms after the onset of the stimuli. In order to test if these data were usable to test our hypothesis, which would rely on subsets of these data, we first decoded the stimulus orientation from the full 600 ms interval. As in all decoding analyses, this was done separately for attend-left and attend-right trials. We could reliably decode the orientation of the attended stimulus from the MEG data (Figure 5) with an accuracy of 70 % (SD = 1.8 %) for attend-left trials, which is higher than expected by chance (T-test for observed versus shuffled condition labels, t(9) = 34.8, p = 6.6 * 10^−11^). The mean accuracy for attend-right trials was also 70% (SD = 1.8 %, t(9) = 34.4, p = 7.3 * 10^−11^). The orientation of unattended trials could reliably be decoded too (unattend-left: accuracy = 70 ± 2.7 %, t(9) = 23.7, p = 2.0 * 10^−9^; unattend-right: accuracy = 69 ± 2.1 %, t(9) = 28.8, p = 3.6 * 10^−10^).

**Figure 5.**
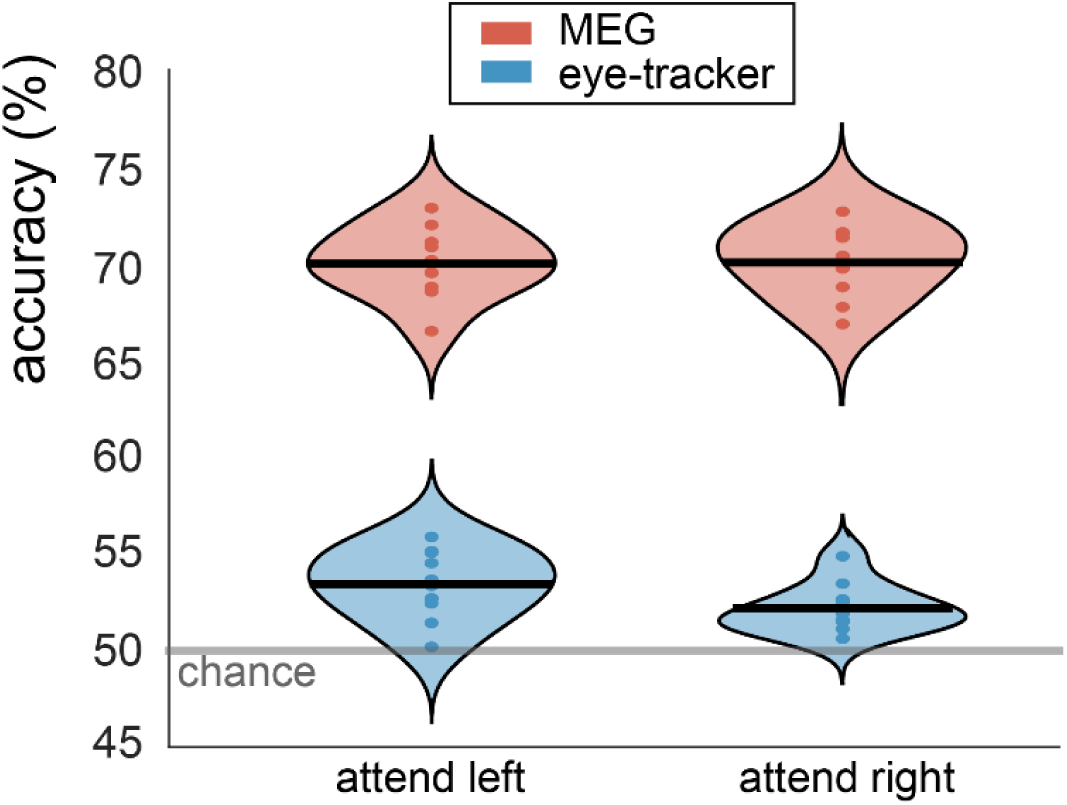
Stimulus information of the attended stimulus can reliably be decoded from MEG data (red) and, to a lesser extent, eye-tracking data (blue). Decoding was done separately for attend-left and attend-right trials. Violin plots display the probability density over subjects. The horizontal line denotes the group mean and the dots denote the individual subject accuracies. The semitransparent gray line shows the empirical chance level.

We ascribe these effects to the difference in cortical processing of gratings with different orientations. However, an alternative explanation could be subtle, yet systematic differences in eye position and/or eye movements between conditions. This is because eye movements affect the MEG signal, and above chance decoding could thus be a trivial result of eye movements (Thielen et al., 2019). Indeed, we were able to decode stimulus orientation with above-chance accuracy from just the eye tracker data (figure 5), but the accuracy was much lower than for the MEG data. The average accuracy was 53 % for attend-left (SD = 1.8 %, t(9) = 6.0, p = 2.1 * 10^−4^) and 52 % for attend-right trials (SD = 1.3 %, t(9) = 5.6, p = 3.4 * 10^−4^). This indicates that there is a bias in gaze depending on the stimulus orientation. We conducted a control analysis in order to exclude the possibility that above chance decoding of MEG data was exclusively a result of either eye movement induced artifacts, or because the visual stimulus ended up in different parts of the retinotopic visual cortex as a consequence of differences in gaze. We trained an SVM to distinguish orientation based on only the eye tracker data and evaluated this at the single-subject level. Next, two pseudo-trials (i.e. average of 5 trial-time points, see 2.3.7), one for each class, with the largest distance from the decision boundary, were removed. This was done separately for each fold. After this, the remaining data were repartitioned into training/test data. This procedure was repeated until the accuracy over ten folds did not surpass chance level 5 times in a row. On average, the percentage of pseudo-trials that had to be removed from the data in order for the accuracy to drop to chance level was 3.3 % (SD = 2.7 %). The same trials were then discarded from the MEG data, which were subsequently used in another decoding analysis. The average decoding accuracy from the MEG data remained at 70 % (SD = 2 %) for both attend-left and attend-right trials, and did not statistically differ from the accuracy level before the removal of pseudo-trials. Table 1 in the Supplemental Information lists the decoding performance at the single-subject level. These results show that even when discarding the trials that are most discriminating based on eye movements, it is still possible to decode stimulus orientation from MEG data with high accuracy. This indicates that successful decoding of stimulus orientation in our data does not rely on differences in gaze or eye movements.

**Table 1.**
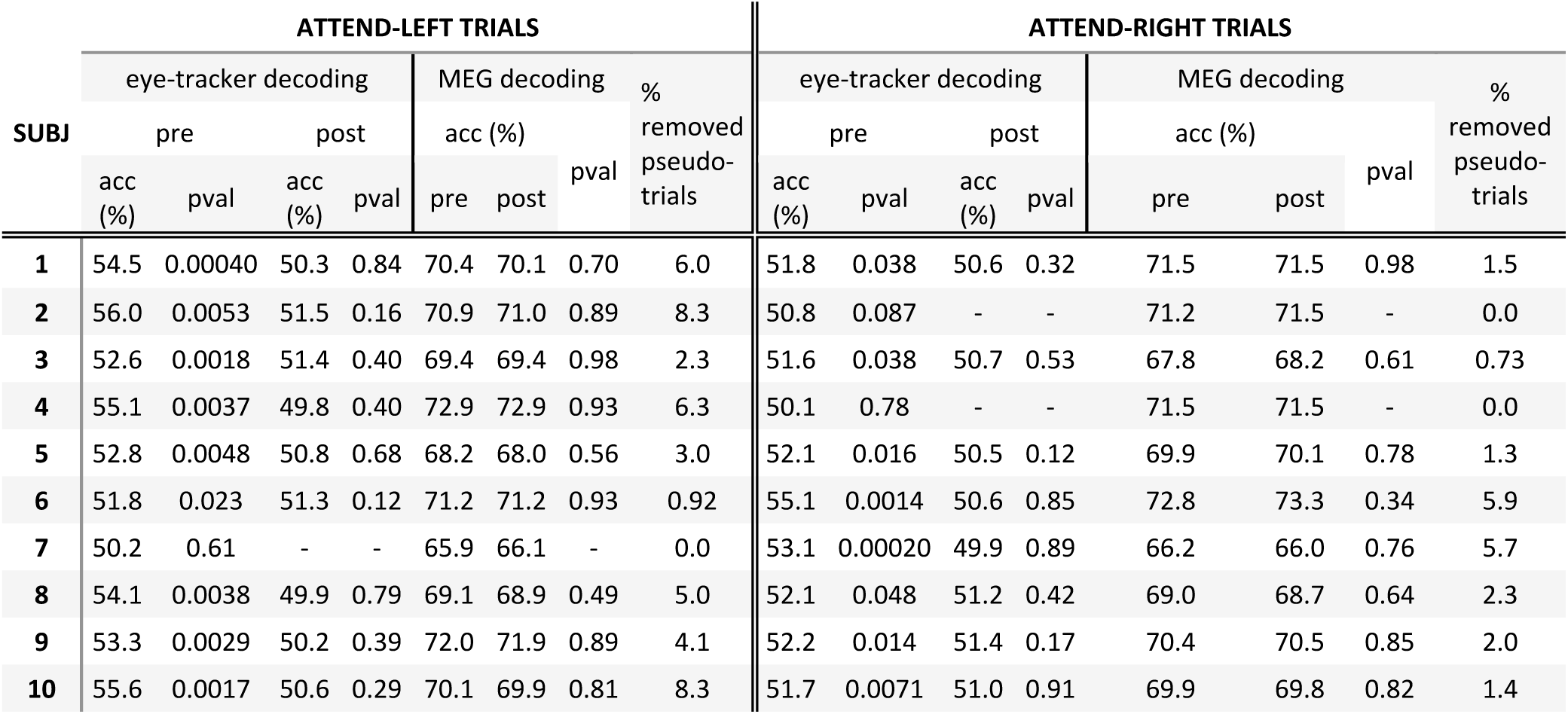
Single subject data from the control analysis on decoding on eye-tracker data. The most discriminative trials were removed from the eye-tracker data until accuracy was at chance level. The same trials were removed from the MEG data and performance with and without removal were compared. The columns in the ‘eye-tracker decoding’-field indicate average accuracy over 10 folds, before and after removal of the selected trials, and the p-value from a T-test of the accuracy relative to the accuracy resulting from shuffled class-labels. The columns in the ‘MEG decoding’-field indicate the accuracy before and after removal of the selected pseudo-trials, and the p-value is the result from a T-test, comparing the accuracy before and after removal of the selected pseudo-trials over folds (i.e. a p-value of >0.05 indicates no difference in MEG decoding accuracy despite removing the pseudo-trials that were furthest away from the decision boundary in the eye-tracking decoding). The last column from in both parts indicate the percentage of pseudo-trials that were removed from the data. Cells are empty in case performance based on eye-tracking data was already at chance level. acc: accuracy; pval: p-value.

### 3.3 Visual representations are not periodically modulated by oscillatory activity in visual cortex

Now that we established that it is possible to decode stimulus orientation information from the MEG signal, we can look into fluctuations in decoding accuracy. Specifically, we hypothesized that decoding accuracy depends on the theta/alpha phase of ongoing activity in relevant brain areas. Specifically, we expected that the phase of brain activity in the regions where the stimulus is actively processed would influence decoding performance. Since gamma synchronization is a proxy for stimulus processing, we used the location of the maximum increase in induced gamma power to provide the frequency-specific phase signal that might modulate decoding performance. All data (i.e. from either attend-left or attend-right trials) was binned according to 18 phase bins, and the orientation of the attended stimulus was decoded from the MEG data separately for each phase bin. This was done independently for frequencies in the theta, alpha, and beta band (4-30 Hz), in steps of 1 Hz. Resulting from this analysis were 18 accuracy scores, one for each phase bin. A cosine function was fit to these scores to estimate the strength of phasic modulation (figure 6A). This was tested against a permutation distribution in which the phases within a trial were shifted randomly, whilst keeping the autocorrelation structure intact (see 2.3.7). Contrary to our hypothesis, the phasic modulation in the observed data was not significantly larger than that of the permutation distribution. The largest effect in attend-left trials was present at 8 Hz, with an average modulation strength of 0.98 % (SD = 0.43 %, p = 0.68, uncorrected). There was also a peak at 11 Hz (mean ± SD = 0.98 ± 0.52 %, p = 0.85, uncorrected, figure 6B). The largest effect in attend-right trials was 0.93 % (SD = 0.38 %, p = 0.85, uncorrected) at 9 Hz (figure 6C). Although the spectral shape of the modulation depth suggests an effect in the alpha band in both conditions, the observed peaks were not statistically significant.

**Figure 6.**
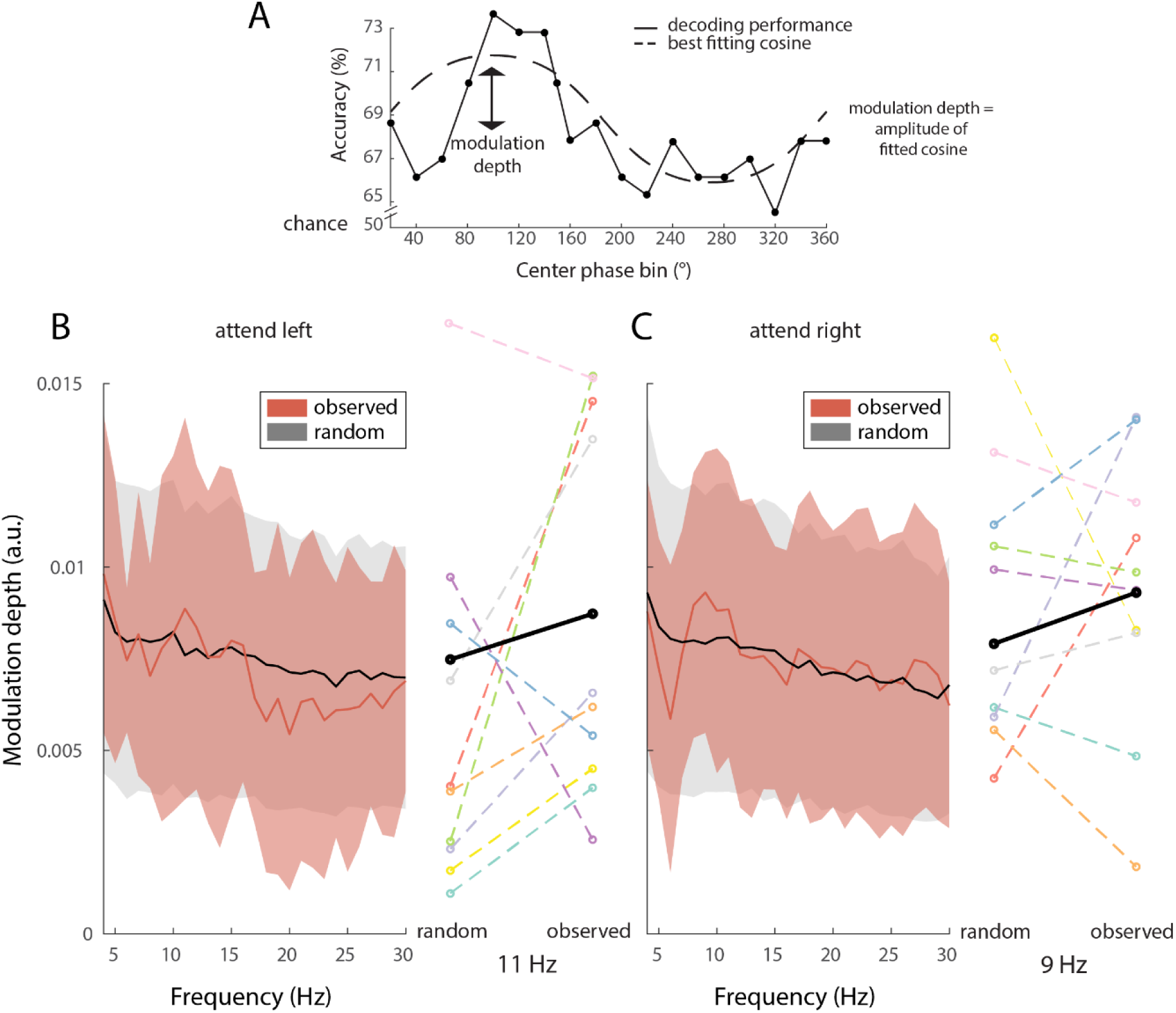
Phasic modulation of decoding performance. A) Schematic of the phasic modulation depth, i.e. the amplitude of a cosine fitted to the decoding performance over phase bins. B and C) Modulation depth as a function of frequency for attend-left (B) and attend-right (C) trials. Observed data in red; random data in grayscale (mean and standard deviation over subjects). Individual effect sizes at the spectral peak are shown on the right of the spectrogram, with the group average in black.

### 3.4 Activity in the fronto-parietal network periodically modulates visual representations

The previous results hinted at an alpha frequency phasic modulation of decoding performance, but the variability over subjects was large and the effects were not statistically significant. A possible reason for this is that the phasic modulation was estimated, based on the signal phase in visual brain areas, whereas the real modulatory signal has a different origin. Some previous reports suggest that the modulatory effect of theta/alpha phase on behavior originates from areas in the fronto-parietal attention network, especially the frontal eye fields (FEF) and parietal cortex (Busch et al., 2009; Helfrich et al., 2018). To investigate this, we performed an additional set of analyses, now using as phase-defining signal source reconstructed activity from anatomically defined cortical parcels from FEF and parietal cortex, resulting in 36 brain parcels (figure 2) in 17 frequency bins (4-20 Hz). We explored whether the phase of the signals from these parcels and frequencies modulated decoding performance. Thus, the analysis strategy used before was now repeated, independently for each parcel and frequency, i.e. the phase time course was estimated on a single parcel and used for subsequent binning in the decoding procedure. We found that phasic modulation was higher than expected by chance in both attend-left and attend-right trials. In attend-left trials, the right FEF most contributed to this difference at 20 Hz (mean ± SD = 1.0 ± 0.44 %, cluster-corrected p = 0.002, figure 7C). Modulation by the right FEF at theta frequency (4 Hz, figure 7A), and by the right parietal cortex in the alpha band (8-11 Hz) were also present (figure 7B). In attend-right trials a cluster in the left FEF at 10-18 Hz mostly contributed to the significant difference (mean ± SD = 1.0 ± 0.49, p<0.001, figure 7E). There were also consistent effects in the left parietal cortex in the theta band (4 Hz, figure 7D) and alpha band (8-12 Hz; figure 7F). The optimal phases for decoding differed between subjects, but were to a large part consistent over parcels and frequencies (see figure S2 and S3). Note that phase polarities are ambiguous: bimodal distributions with a 180 degree phase difference between adjacent parcels can occur due the polarity ambiguity in the principal component analysis (PCA) in the construction of parcel time courses (see Methods 2.3.5). The subjects with the largest modulation effects (figure 7) seem to have a more consistent optimal phase within neighboring parcels and frequencies (for example, compare the modulation strength of subjects S1 (“good” subject) and S4 (“bad” subject) in figure 7D and their phase distributions in figure S3A), which adds to the credibility of the modulation effect.

**Figure 7.**
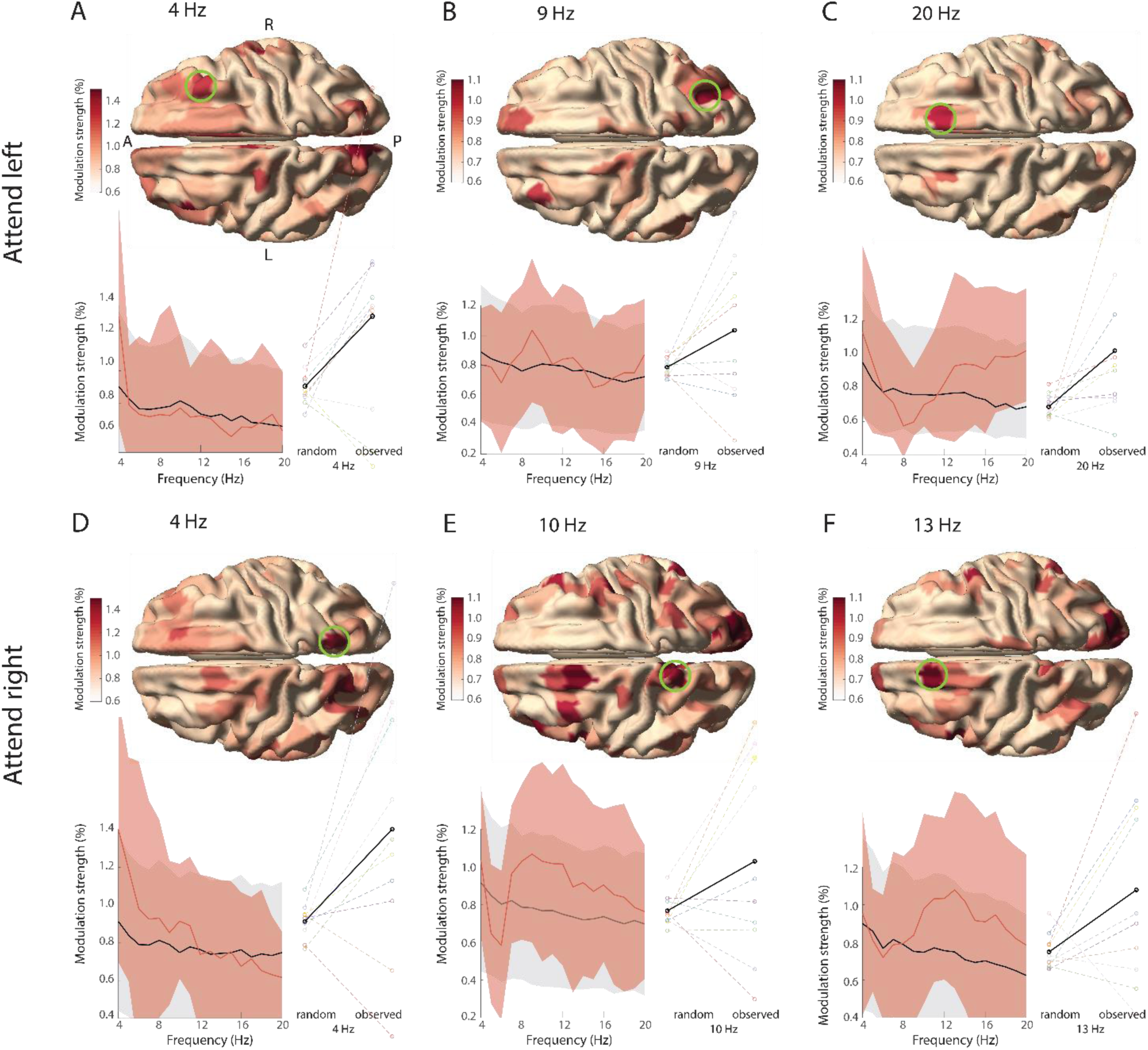
Phasic modulation of decoding performance in the fronto-parietal network. The phasic modulation in attend-left trials (A-C), and in attend-right trials (D-F) is shown for all parcels, but only the parcels in the ROIs were tested statistically (see figure 2) as described in 2.4.2. In A the anterior (A) to posterior (P) axis and left (L) and right (R) hemispheres are denoted. Spectrograms show the group-average modulation strength in the selected parcel in the green circle (red) and the average expected by chance (black). Shading reflects S.D. across subjects. Individual effect sizes at the spectral peak are shown on the right of the spectrogram, with the group-average in black.

### 3.5 Left fronto-parietal areas might modulate reaction times

Having established that the phase of ongoing theta/alpha activity in FEF and parietal cortex modulates the strength of cortical stimulus representation, we next investigated whether the phase of these signals modulates behavioral performance as well. This has been reported in other studies (Helfrich et al., 2018, 2018). Specifically, we tested whether the phase in these areas at the moment of the stimulus change (i.e. to which subjects had to respond) affected reaction times. A cluster-based permutation test within the same ROIs as before revealed no phasic modulation of reaction times: for attend-left trials, p = 0.23 (corrected); for attend-right trials, p = 0.13 (corrected). However, some interesting trends were observed. In attend-left trials, the largest effect was present in left FEF parcels at the lower end of the alpha band (8 Hz), with a mean modulation strength over subjects of 1.1 % (SD = 0.55 %, p = 0.53, uncorrected), with similar trends in right and left parietal cortex (see figure S1). In attend-right trials, the largest effect was also present in the left FEF at the higher end of the alpha band (13 Hz), with a mean modulation strength over subjects of 1.0 % (SD = 0.63 %, p = 0.13, uncorrected).

Our results show that stimulus information can be decoded with a higher accuracy at particular phases of the theta/alpha rhythm in the fronto-parietal network contralateral to the attended hemifield. This is consistent with the idea that visual perception is not continuous but to some extent discrete. The behavioral relevance of the phase in this network could not be confirmed statistically, but trends suggested that there might be a modulatory effect on reaction times.

**Figure 8.**
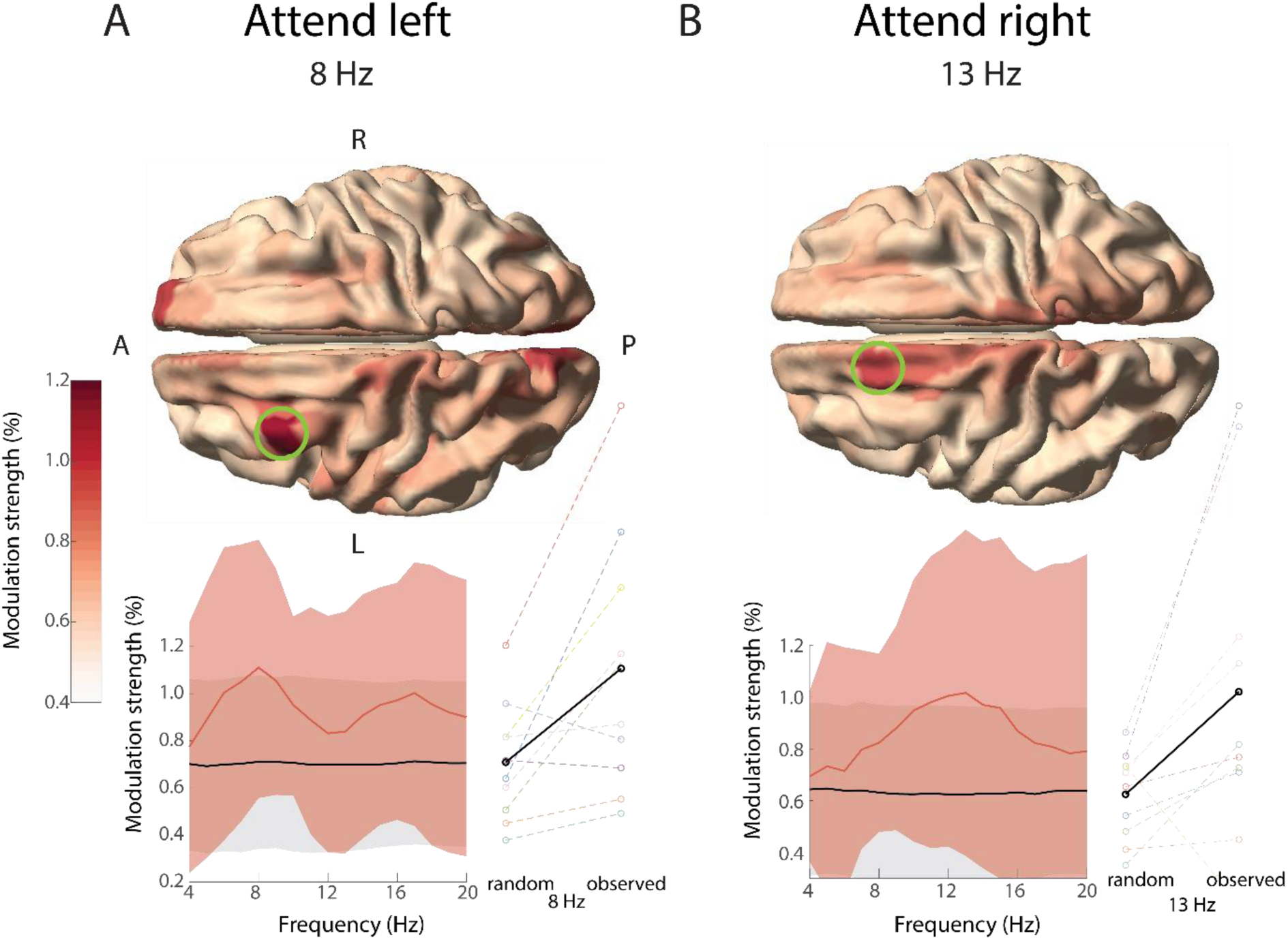
Similar to figure 7, but now showing the phasic modulation of reaction times. The group average modulation strength peaks in frontal and parietal areas in the alpha range, with a dominance of the left hemisphere in both attend-left and attend-right trials, but was not significant.

## 4. Discussion

In this study, we investigated the presence of rhythmicity in visual representations during sustained spatial attention. Psychophysical and neural evidence suggest that (visual) perception is not continuous but discrete, as a consequence of discrete rhythmic attentional sampling. We reasoned that if this is the case, neural representations are also expected to fluctuate, i.e. the strength of a neural representation should depend on the phase of ongoing activity. This was tested by decoding stimulus orientation from human brain signals, as a function of phase of the ongoing activity. We found that oscillatory activity in visual cortex does not phasically modulate decoding accuracy. However, the frontal eye fields (FEF) and parietal cortex contralateral to the attended stimulus did phasically modulate decoding accuracy in the theta and alpha bands. The phasic modulation of visual representations is in line with discrete perceptual sampling during attention, though it is unclear at which step of visual processing it occurs. The behavioral relevance of the phase of ongoing activity in this network was not confirmed, but trends in the phasic modulation of reaction times were in line with such modulatory effect.

The phasic modulation of decoding performance was mostly present in the theta and alpha bands. These effects were band-limited and in no occasion did the effect extend to the entire investigated frequency range. This suggests that the above chance modulation effects were not a consequence of a bias in the permutation test or of other confounding effects. Effects in these frequency bands are also in line with the perceptual sampling literature. Busch et al. (2009) showed that the phase of the frontal theta/alpha rhythm (6-12 Hz) predicts whether a threshold stimulus is perceived or not.

Similarly, Fiebelkorn et al. (2018) and Helfrich et al. (2018) showed that perceptual outcome depends on the theta phase in the fronto-parietal network (including FEF and parietal areas), and that this is accompanied by increases in cortical excitability during ‘good’ theta phases. It is not yet clear whether phasic modulation in the theta and alpha band reflect the same or different neural processes, but evidence from phasic modulation of behavioral performance suggests the presence of either rhythm might be task-dependent and rely on the number of objects that have to be monitored (Fiebelkorn et al., 2013; Holcombe and Chen, 2013; Landau and Fries, 2012). This would mean that the theta and alpha band effects are not necessarily related to different processes per se, but rather to temporal limits of attentional selection. The spatial locations of the theta/alpha effects slightly differed between attend-left and attend-right trials. In attend-left trials, a theta modulation effect was present in the FEF and an alpha modulation effect in parietal cortex. In attend-right trials, the theta modulation effect was strongest in parietal cortex and the alpha modulation effect in parietal cortex and FEF. A previous report on the neural correlates of rhythmicity in behavioral performance also showed variability in the topographical origins of the phasic modulation (Helfrich et al., 2018). Possibly, the entire fronto-parietal network is involved in the rhythmic nature of sustained attention, but different parts of the network have different effects. Fiebelkorn et al. (2018) suggest the FEF might specifically be involved in (the attenuation of) exploration of space, while the parietal cortex modulates processing at the attended location. In line with this, Gaillard et al. (2020) used a cued target-detection task in macaques and showed that the attentional spotlight explores space rhythmically, by decoding the location of the attentional spotlight from the prefrontal cortex (PFC). Future research should thus disentangle the different contributions of the frontal and parietal cortex in rhythmic sampling. For example, one could conceive of an experiment in which the setup of the Gaillard et al. are combined with the approach in the current study. If it is possible to decode stimulus information and the position of the attentional spotlight on the same trial, we would expect both to be modulated by theta/alpha phase, but they would rely on different origins (e.g. parietal versus prefrontal cortex) or have different optimal phases.

We did not observe a phase dependent modulatory effect when using visual cortical activity as the phase delivering signal, which is at odds with the previous literature. Landau et al. (2015) found that that the sampling of two behaviorally relevant stationary grating stimuli occurred at 4 Hz each and that this modulation was present at cortical locations with maximal visually induced gamma-band activity. The current study used a very similar experimental paradigm (albeit with drifting instead of stationary gratings, and a different attentional manipulation), and it is likely that the distributed activity in early visual areas played a major part in successfully decoding stimulus orientation. Primary visual cortex is most sensitive to local contrast differences like oriented lines (Hubel and Wiesel, 1962), and empirical and modelling work from Cichy et al. (2015) confirmed it is likely that orientation information is decoded from this region with MEG (Stokes et al., 2015). The discrepancy in the locus of the phasic modulatory signal requires further investigation. Especially the question whether perceptual sampling in visual areas is always under control of the fronto-parietal attention network, or whether it can be controlled locally. Some evidence for the former comes from Popov et al. (2017), who found that processing in visual cortex (as measured by gamma power) was a function of the attentional modulation of posterior alpha power, while the latter was Granger caused by the FEF. Activity in the FEF itself was not modulated by attention. Consequently, in the current study as well, visual processing might be modulated by the FEF without clear changes in spectral power in that region (figure 2). There are a number of potential reasons why the relationship between theta/alpha phase and decoding accuracy was not observed from a source in visual cortex. It could be that the induced visual cortical activity reflects multiple generators of oscillatory activity with heterogeneous phase relationships (Maris et al., 2016), thus prohibiting the estimated phase values to actually reflect the physiologically relevant phase. This possible explanation is supported by the presence of multiple generators of posterior alpha with different sensitivity to different visual locations (Popov et al., 2019). Another option is that visual representations might not be modulated in early visual cortex, but in downstream visual areas. This is in line with evidence from fMRI studies: Sprague and Serences (2013) showed that attention increases the amplitude of stimulus representations especially in higher visual areas. Moreover, orientation-selective representations are not only present in visual areas, but also in parietal and frontal areas including the FEF, and these areas are all modulated by attention (Ester et al., 2016).

We did not find any significant modulatory effects of neural oscillatory phase on behavior, in contrast earlier findings. Studies relating behavioral performance to the phase of ongoing activity typically use binary performance measures, e.g. hits versus misses (Busch et al., 2009; Busch and VanRullen, 2010; Fiebelkorn et al., 2018; Helfrich et al., 2018). Perhaps modulatory effects in these performance measures can be detected with higher sensitivity than a continuous variable like reaction time. Note that the strongest (non-significant) effects in behavioral performance were present in the FEF and parietal cortex, in the alpha band, which is in line with the origins of the modulation effects in decoding performance. It is therefore plausible that the phasic modulation of visual representations and behavioral performance are a consequence of the same attentional process, but this should be confirmed empirically.

Multivariate pattern analysis (MVPA) has been used before to investigate how neural representations might be related to the alpha rhythm (Foster et al., 2017b, 2017a, 2016; Samaha et al., 2016; van Moorselaar et al., 2018). In those studies, neural representations of the locus of spatial attention were decoded from the topographical distribution of spectral power. The location could be decoded specifically from alpha band power, both during attention selection and working memory (WM) maintenance, even when spatial position was irrelevant to the WM task. This shows that there is a link between the spatial distribution of alpha band activity and the representation of space, and one might ask whether and how this is related to the modulation of neural representations by alpha phase in the current study. We also used a covert spatial attention task, and the attentional modulation index (AMI) showed a lateralization in posterior alpha power as a function of direction of spatial attention (figure 3), which confirms a link between alpha activity and the locus of spatial attention. However, the stimulus feature (i.e. orientation from the *attended* stimulus) we decoded from the MEG data was independent from its spatial location (i.e. only the orientation of the attended stimuli were compared when the attended stimulus was in the same hemifield). In fact, all attend-left and all attend-right trials were analyzed separately in order to ensure that decoding of stimulus orientation was not confounded by spatial attention. Furthermore, the topographical distribution of alpha activity did not play any part in the decoding analysis, since the phase binning was based on a single and the same alpha source for both orientations. Therefore, it is unlikely that the representation of space in a signature pattern of alpha band activity contributed to, or confounded, the phasic modulation of neural representations.

Recently, a number of studies have demonstrated confounding effects of eye movements on decoding performance from M/EEG data (Mostert et al., 2018; Quax et al., 2019; Thielen et al., 2019). Because eye movements affect the MEG signals, a systematic difference in gaze position between conditions could result in above-chance decoding performance independent of differences in the true underlying neural representation. We found that it is possible to decode stimulus orientation reliably from the eye tracker data only, which reinforces this concern. In order to make sure that the MEG decoding was not a trivial result of eye movements or gaze position, we investigated whether orientation could still be decoded from the MEG data after removal of the trials that could distinguish orientation based on gaze. Removal of these trials resulted in chance level accuracy based on eye-tracker data, but did not affect the accuracy based on MEG data, which remained highly above chance level. Another concern could be a contamination of the decoding performance by evoked activity. Temporal MEG decoding performance is highest at the peak of evoked activity Cichy et al., 2015; Pantazis et al., 2018; Ramkumar et al., 2013). At the same time, these peaks trivially have a consistent phase over trials, which could lead to a trivial phasic modulation in decoding performance. We ensured that evoked activity did not affect estimates of phasic modulation effects in decoding by only using data for the decoding analysis after offset of the evoked activity, which was not phase-locked (data not shown).

Overall, this exploratory study shows that visual representations are modulated by the phase of ongoing activity. In particular, it indicates that the representation of visual input is not constant over time, but depends on the phase of the theta/alpha rhythm in the fronto-parietal attention network. Further research should investigate the potentially differential roles of the frontal and parietal parts of this network on the one hand, and frequencies in the theta and alpha band on the other hand.

## 6. Supplemental information

**Figure S1.**
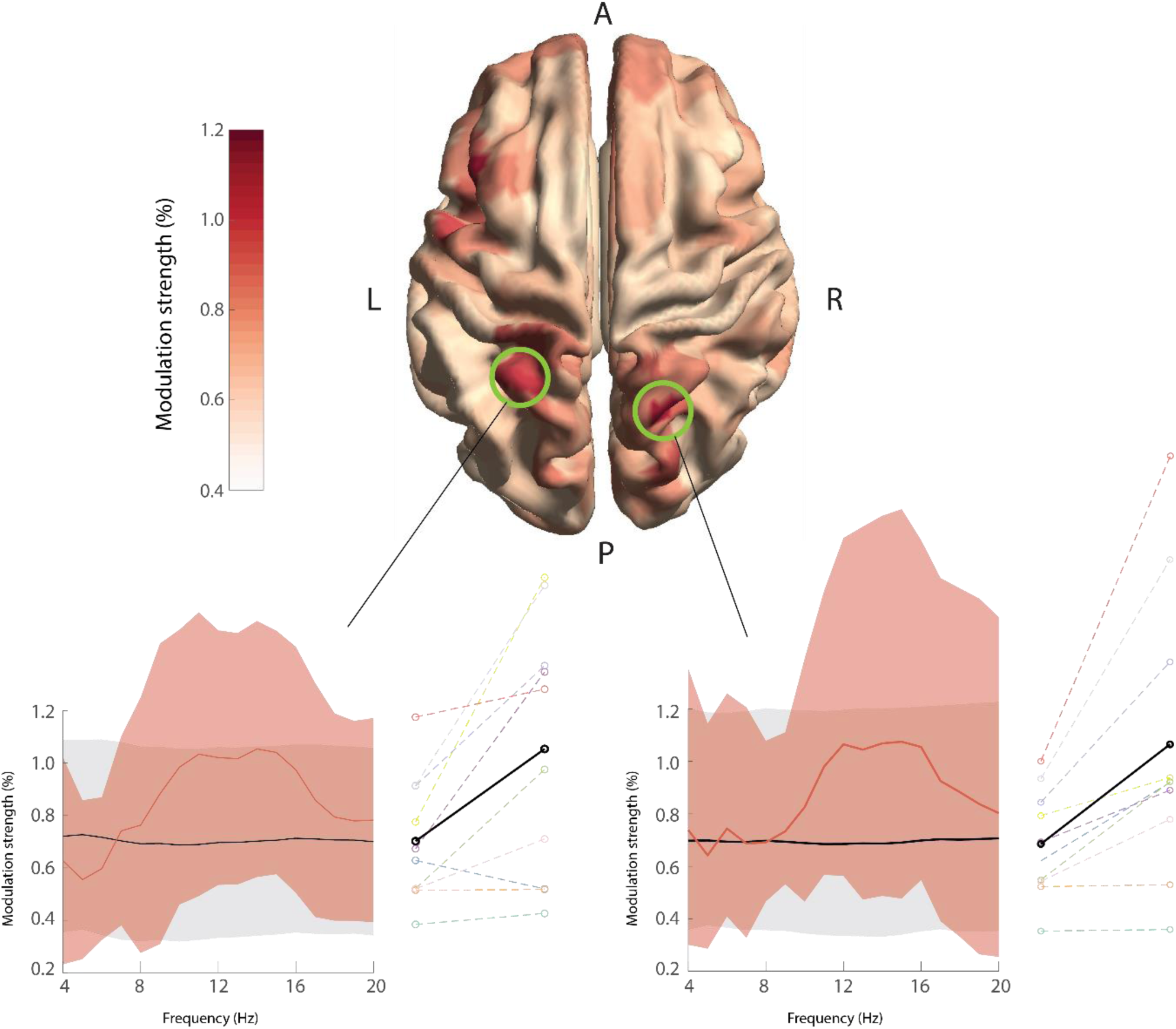
Phasic modulation of behavior in attend-left trials. Similar to figure 8, it displays a peak in modulation strength in the alpha band in left and right parietal cortex.

**Figure S2.**
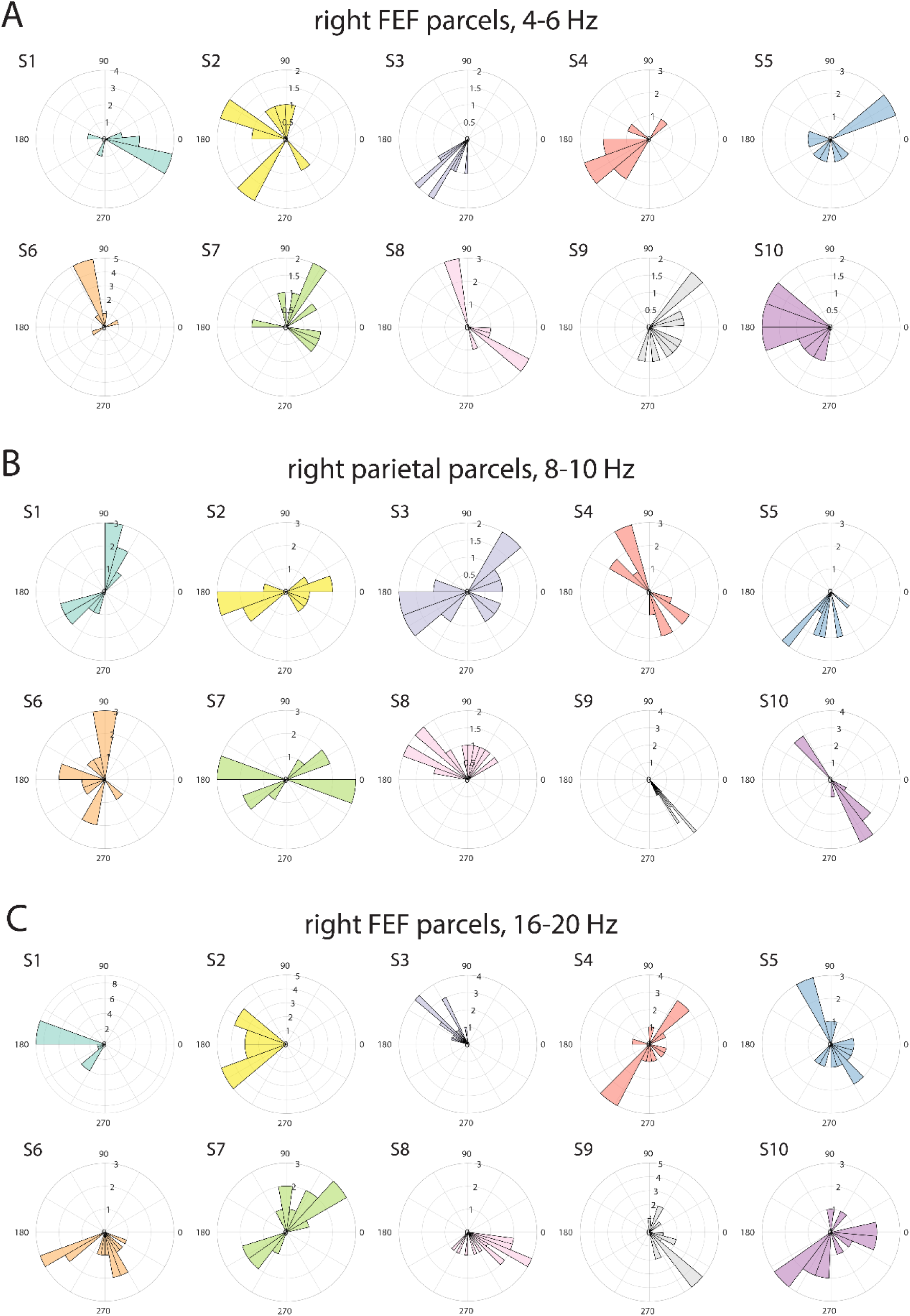
Optimal phase for decoding in attend-left trials. Histograms of optimal phases over parcels and frequencies that contain the strongest phasic modulation effect. Each subplot corresponds to the subplot in figure 7A-C in the same order and subject’s colors are matched with the right most panel in those subplots. Note that phase polarities are ambiguous (see main text): bimodal distributions with a 180 degree difference suggest similar optimal phases of frequencies and parcels.

**Figure S3.**
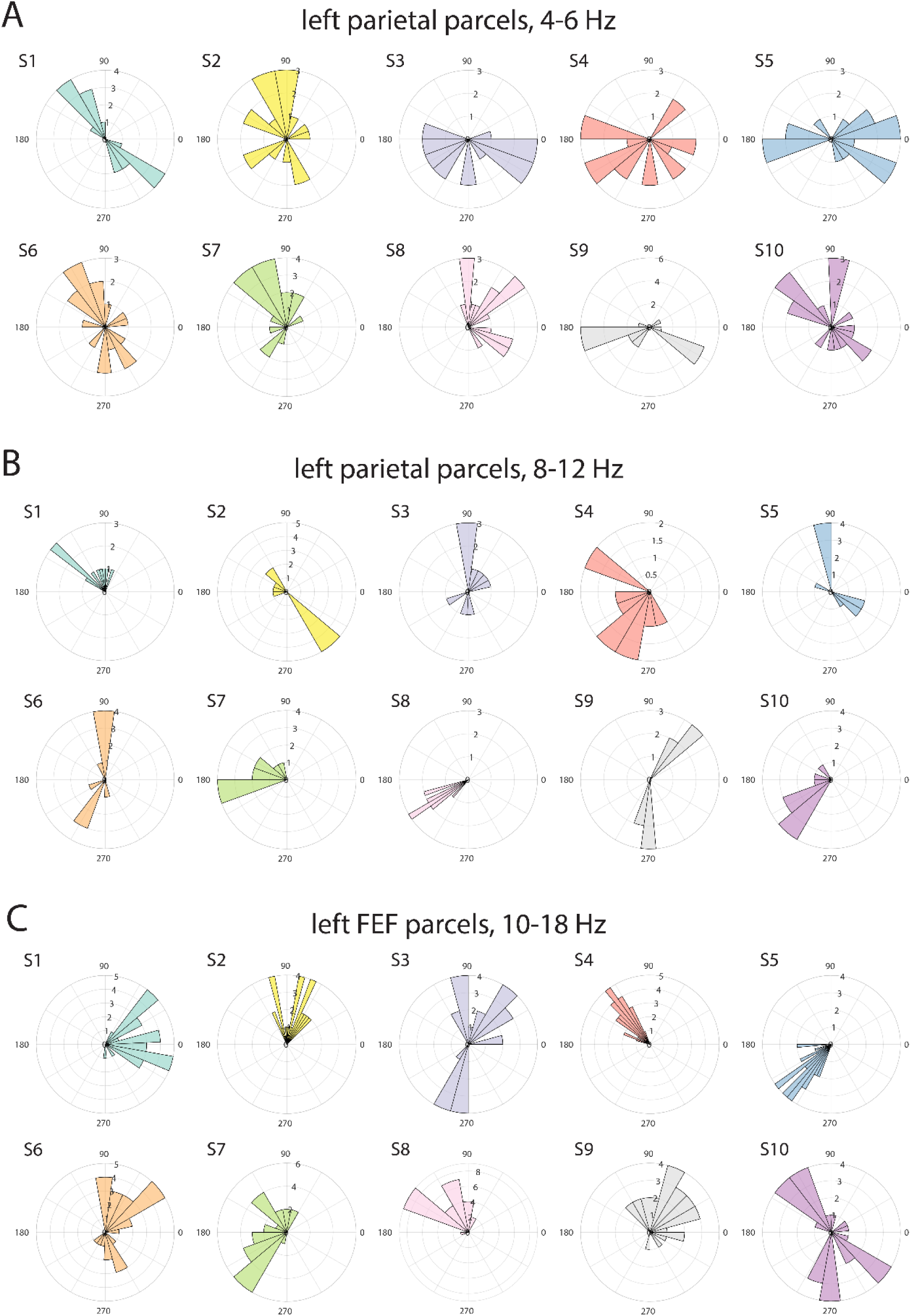
Optimal phase for decoding in attend-right trials. Histograms of optimal phases over parcels and frequencies that contain the strongest phasic modulation effect. Each subplot corresponds to the subplot in figure 7D-F in the same order, and subject’s colors are matched with the right most panel those subplots. Note that phase polarities are ambiguous (see main text): bimodal distributions with a 180 degree difference suggest similar optimal phases of frequencies and parcels.

